# Ambient light color does not detectably change the transient, flight-specific DEET response of *Aedes aegypti*

**DOI:** 10.64898/2026.10.23.761422

**Authors:** Serene Aryal, Amer Dawoud

## Abstract

We asked whether ambient light color changes how *Aedes aegypti* respond to the repellent DEET. We reanalyzed an unpublished image dataset from the chamber system of Carver [2018]: 12 recordings of about 20 mosquitoes each, under six light colors (blue, green, orange, red, white, yellow), with one recording per color in each of two sessions. Each recording had about 10 minutes with bait only and then about 10 minutes with bait plus DEET. We report two robust findings and one negative result. First, the DEET response is short-lived. Pooled across all 12 recordings, flight motion fell by about half in the minute of DEET onset and recovered within about three minutes (ratio of means 1.99, exact sign-flip one-sided *p* = 0.058); over the full halves there was no net difference (ratio of means 1.01, *p* = 0.49). Second, the drop is a loss of flight, not a loss of detection. A separate presence readout, which also counts resting mosquitoes, stayed almost flat (5.6 to 5.1 detections per frame) while flight fell from 0.51 to 0.35 events per frame pair. The chamber is closed, so presence by itself is not a finding; it shows that the animals remained detectable while flight collapsed, and the recovery of flight excludes mortality. Third, we could not show that color changes the response. A detectability control ruled out an imaging artifact. A baseline-activity control then showed that the apparent color ranking follows how much the mosquitoes flew before DEET (Spearman +0.83). Under a null with no color effect, 1.5 of 6 colors pass the cross-session concordance bar by chance. After adjusting for baseline activity and for selection among six colors, no color stood out (*p* = 0.47). Ratio-of-halves suppression measures in chamber assays are confounded with baseline activity, and prospective studies should match activity across conditions or use time-matched within-recording contrasts.

## 1 Introduction

Female mosquitoes combine smell, vision, and heat to find hosts [van Breugel et al., 2015, Vinauger et al., 2019, Zermoglio et al., 2017, Chandel et al., 2024]. Spatial repellents act on this behavior rather than by killing: they can drive movement away from a source, block host detection, or stop feeding [World Health Organization, 2013, Achee et al., 2012]. DEET is the reference repellent for behavioral work, but its effect is not uniform; some *Aedes aegypti* become behaviorally insensitive to it [Stanczyk et al., 2010]. How the response changes over time, and whether an initial response lasts, is measured less often than its average size.

Vision matters in mosquito sensory biology. Mosquitoes link odor plumes to visual and thermal targets [van Breugel et al., 2015], combine odor and vision in host-seeking flight [Vinauger et al., 2019], and prefer some visible colors when odor is present [Alonso San Alberto et al., 2022], although human odor can dominate landing decisions [Sumner and Cardé, 2022]. It is therefore reasonable to ask whether ambient light color changes a repellent response. None of these studies tested DEET or light, so the link is a hypothesis, not an expectation.

Carver’s 2018 Master’s thesis built a chamber system with gas sensors and camera-based motion counting to measure DEET behavior, and noted possible habituation at higher doses [Carver, 2018]. The same paradigm was also run under six LED colors, but those recordings were never analyzed. Here we analyze them. The design has one recording per color per session, so a color-by-DEET interaction cannot be tested, and a significance test at that replication would be false precision. We therefore ask what these recordings can show and what they cannot. Two controls frame the answer: one rules out an imaging artifact from a monochrome camera under colored light; the other tests whether the remaining color differences can be separated from baseline activity. They cannot, and we report the color comparison as an instructive negative result.

## 2 Methods

### 2.1 Recordings and dataset

We used archived monochrome image recordings from the chamber system of Carver [2018]. The chamber is a closed glass cylinder (about 20.75 in long, 12 in outer diameter, 40.59 L), imaged by a Baumer LXG-120M monochrome camera at 4096 *×* 3072 pixels and 19 frames per second, with about 20 *Ae. aegypti* per recording. Each recording had a bait-only part and then a bait-plus-DEET part, with the switch at frame 11400 (about 10 min). The DEET dose was 1 mL of 95% DEET (about 24 ppm nominal in the chamber volume, per the thesis). The chamber is sealed, so the mosquitoes could not enter or leave during a recording; this matters for the presence measure below. Light color was blue, green, orange, red, white, or yellow. Calibrated spectral peaks were not in the archived metadata, so we report results by color category, not by wavelength.

The dataset had 12 recordings: one per color in each of two sessions (`Color Test 1` and `Color Test 2`), which serve as replicate sets. An automated inventory confirmed all 12 recordings, the DEET-onset frame in each, and no missing frame indices; each recording gave 22,800 usable frames (0–11399 bait only; 11400–22799 bait plus DEET). A separate 100-frame `Ref` directory in session 2 was not used, and a single empty trailing image file in a few recordings was skipped. The chamber was re-set between sessions, so we calibrated the region of interest (ROI) separately for each session and checked it against frame overlays. The pipeline retried transient image-read failures and rejected any recording that did not yield its full set of frame pairs.

### 2.2 Counting flight (motion events)

We followed the thesis method rather than identity-preserving tracking. For each pair of consecutive frames, the pipeline converted the images to grayscale, downscaled them by a factor of 0.35, and took the absolute difference. It thresholded the difference at 12 gray levels, applied erosion (3 *×* 3) and then dilation (5 *×* 5), and kept only the chamber ROI. It retained external contours with area 4 to 250 px (at the downscaled size) and assigned each contour centroid to one of seven zones along the long axis of the chamber. For each frame pair it recorded the number of motion contours, the total motion area, and per-zone counts. It also produced one-minute (60 s) binned traces.

This readout counts motion events, not mosquitoes. A resting mosquito is not counted, and one flying mosquito adds an event in each frame pair in which it moves. We treat it as a relative index of flight activity.

### 2.3 Counting presence (occupancy)

Because the motion readout ignores resting mosquitoes, we built a second, independent readout of presence. In each frame we found dark blobs with a morphological black-hat filter (21 *×* 21-px elliptical kernel), which is larger than a mosquito but smaller than the illumination gradients, so it finds resting and flying mosquitoes alike. Fixed dark features (glass edges, hardware) recur at the same place in every frame. We therefore removed any detection that recurred at the same location in at least 70% of sampled frames. Mosquitoes move, so they survive this filter. Detection used the same per-session ROI and 0.35 resize as the motion pipeline, a threshold of 14 gray levels, and one frame in every 57. The per-frame detection count is the occupancy estimate. Overlays confirmed that detections fell on individual mosquitoes, typically 5–10 per frame for about 20 animals, so the count is a lower bound. Mosquitoes settle after they are introduced, so presence declines over the first minutes regardless of DEET, and the full-half occupancy comparison is confounded by this trend. We therefore read occupancy only through the time-matched onset window and the overall time course, never as a per-color effect size.

### 2.4 Two controls

#### Detectability control

The camera is monochrome and the light color varied, so a per-color difference in measured activity could be an imaging artifact. For 200 evenly spaced frame pairs per recording, using the same detection steps as the main pipeline, we measured: (i) scene brightness (mean gray level in the ROI); (ii) the background noise floor (frame-to-frame absolute difference at ROI pixels outside any detected blob, summarized by its 95th and 99th percentiles and the fraction of background pixels above the threshold); (iii) the motion signal (peak absolute difference inside each detected blob) and its margin above the threshold; and (iv) the signal-to-noise ratio (signal median divided by noise 95th percentile). We then swept the threshold over 6, 8, 10, 12, 16, 20, and 24 gray levels and asked whether the rank order of the six colors changed (Spearman correlation of each ranking against the ranking at 12). Finally we correlated each color’s baseline motion rate with its brightness, noise floor, and signal-to-noise ratio. If detection drove the ranking, the noise floor would approach the threshold, the signal margin would be small, the ranking would change with the threshold, and activity would track brightness.

#### Baseline-activity control

The detectability control addresses an imaging artifact; a statistical artifact can survive it. The sustained effect measure is a ratio of means, bait-only rate divided by bait-plus-DEET rate, and the bait-only rate is also where the colors differ most. Two problems follow. The baseline rate is the numerator of the ratio, so the two are coupled by construction (the spurious correlation of ratios described by Pearson [1897]). And a color under which the mosquitoes flew very little has little flight for DEET to remove, so its ratio stays near one whatever DEET does. Either route can create a color ranking with no color-specific effect.

We tested three things. (i) The association: the Spearman correlation across the six colors between baseline flight rate and sustained suppression, with an exact permutation *p*-value over all 6! relabelings. (ii) How often the observed pattern appears when DEET has the same effect in every color. In this null each recording keeps its own baseline level and dispersion: both halves are resampled with replacement from its own bait-only minute bins. DEET then acts as one multiplier common to all recordings (0.992, from the pooled data) by binomial thinning. We resampled minute bins, not frame pairs, because flight is bursty within a minute; a half-recording holds about ten quasi-independent units, not 1140. From 20,000 draws we recorded how often the cross-session concordance bar is passed by chance. We also recorded how often the best of six colors reaches the largest observed suppression ratio while concordant, because we picked the leading color after seeing the data. (iii) Whether color explains anything beyond baseline: we regressed recording-level log_2_ suppression on baseline rate, permuted color labels within session (20,000 permutations, each color keeping one recording per session), and asked whether any color’s mean residual was extreme.

### 2.5 Per-color summaries, contrasts, and statistics

Each color received a fingerprint: its minute-by-minute trajectory; the acute drop (fractional fall in motion in minutes 10–11 relative to the pre-onset baseline of minutes 7–9); the recovery (plateau in minutes 13–19 as a fraction of that baseline); the sustained effect (full bait-only half versus full bait-plus-DEET half); and the share of activity in each of the seven zones. Every color-level value is a ratio of means that pools the raw event counts of both same-color recordings; we did not average per-recording ratios, which are unstable at near-zero counts. The two sessions serve to check concordance, that is, whether both agree in direction. Colors with pre-onset baseline activity below 0.15 events per frame pair are flagged and read from the trajectory, because their ratios rest on near-zero denominators. The zone layout (zone 1 near the bait, the DEET source and peak vapor near zones 2–3, zone 7 the far, low-DEET end) comes from the thesis protocol and was confirmed by the original investigator. We did not re-derive the zone-to-chamber mapping, so spatial statements are treated as suggestive.

We computed two contrasts. The *sustained* contrast sampled every tenth frame (about 2,280 frame pairs per recording, about 1,140 per condition) and compared the full halves. The *acute* contrast sampled every other frame in a window around onset (frames 9120–13680, about 2 min on each side) and compared the two minutes before onset with the two minutes after; adjacent minutes minimize slow within-recording trends.

The primary outcome was mean motion events per frame pair in each condition, per recording. The primary effect measure was the recording-level raw difference (bait-only minus bait-plus-DEET; positive means less motion after DEET). As a relative effect we report the ratio of means (summed bait-only motion divided by summed bait-plus-DEET motion, with a recording-level bootstrap interval). All inference was at the recording level (*n* = 12; the unit is one cage of about 20 mosquitoes), using an exact sign-flip permutation test on raw differences and an exact sign test. We avoided a parametric mixed model, which would be falsely precise with two recordings per color. Between-color heterogeneity was assessed with an exploratory permutation ANOVA, with *η*^2^ compared against its small-sample null (*k −* 1)*/*(*N −* 1) = 0.45 for *k* = 6 colors and *N* = 12 recordings. No multiplicity correction was applied, so all color-specific statements are exploratory. The full pipeline and its outputs are released with the manuscript.

## 3 Results

### 3.1 Detection is the same in every color

We first asked whether the monochrome camera under colored light could create a per-color ranking by itself (Table S1; Figure S1). It could not. Scene brightness varied little (mean ROI gray level 91– 101 of 255). The background noise floor was 1–2 gray levels in every color, far below the threshold of 12, and the fraction of background pixels above the threshold was *≤* 4 *×* 10^*−*6^. The motion signal cleared the threshold by +16 to +47 gray levels in every color, so detection is limited by signal, not by the threshold. Sweeping the threshold from 6 to 24 left the color ranking essentially fixed (Spearman *≥* 0.94 against threshold 12 across the range 8–16). Baseline activity did not track brightness (Spearman *−* 0.26) or the noise floor (*−* 0.15); the brightest color, yellow, was the least active, and the most active, white, had intermediate brightness. Activity did correlate with signal-to-noise ratio (Spearman +0.66), but this is not significant at *n* = 6 (critical |*ρ*| *≈* 0.89) and is expected, because mosquitoes that fly more make larger motion blobs. The activity differences across colors are therefore behavioral, not imaging artifacts.

### 3.2 The DEET response is acute and transient

Pooled across all 12 recordings (Figure 1), mean motion fell from about 0.55 events per frame pair in the two minutes before DEET to 0.19 in the onset minute. It recovered to about 0.51 within about three minutes. The acute contrast confirmed this drop. Motion in the two minutes after onset was about half that before (ratio of means 1.99, 95% CI 1.00–4.52; mean raw difference 0.272 events per frame pair; exact sign-flip one-sided *p* = 0.058, two-sided *p* = 0.116; 7 of 11 nonzero recordings suppressed). This is suggestive but not conventionally decisive. The sustained contrast showed no net difference between the full halves (ratio of means 1.01, 95% CI 0.58–1.64; one-sided *p* = 0.49; 5 suppressed, 7 increased; Table 1; per-recording values in Table S3). Excluding the low-signal recordings changed neither conclusion (acute *p* = 0.078, *n* = 7; sustained *p* = 0.476, *n* = 10). The DEET-linked drop in flight is therefore transient: it is concentrated at onset and does not persist. This fits rapid behavioral habituation, which the original thesis also raised [Carver, 2018].

**Table 1:** Overall DEET effect, recording-level inference (*n* = 12). The acute contrast uses the peri-onset window (frames 9120–13680); the sustained contrast uses the full halves. Color-heterogeneity rows are exploratory; compare *η*^2^ with its small-sample null of 0.45. The sensitivity row repeats the primary test without the low-signal recordings (five in the acute pass, two in the sustained pass).

| Statistic | Acute (peri-onset) | Sustained (full halves) |
| --- | --- | --- |
| Ratio of means (bait-only / bait+DEET) | 1.99 | 1.01 |
| 95% bootstrap CI | 1.00–4.52 | 0.58–1.64 |
| Exact sign-flip $p$, one-sided | 0.058 | 0.49 |
| Exact sign-flip $p$, two-sided | 0.116 | 0.98 |
| one-sided, excluding low-signal recordings | 0.078 ($n=7$) | 0.476 ($n=10$) |
| Sign test (suppressed / increased) | 7 / 4 | 5 / 7 |
| Permutation ANOVA $p$ (color, exploratory) | 0.10 | 0.18 |
| Permutation ANOVA $\eta^2$ (null 0.45) | 0.79 | 0.64 |

**Figure 1:**
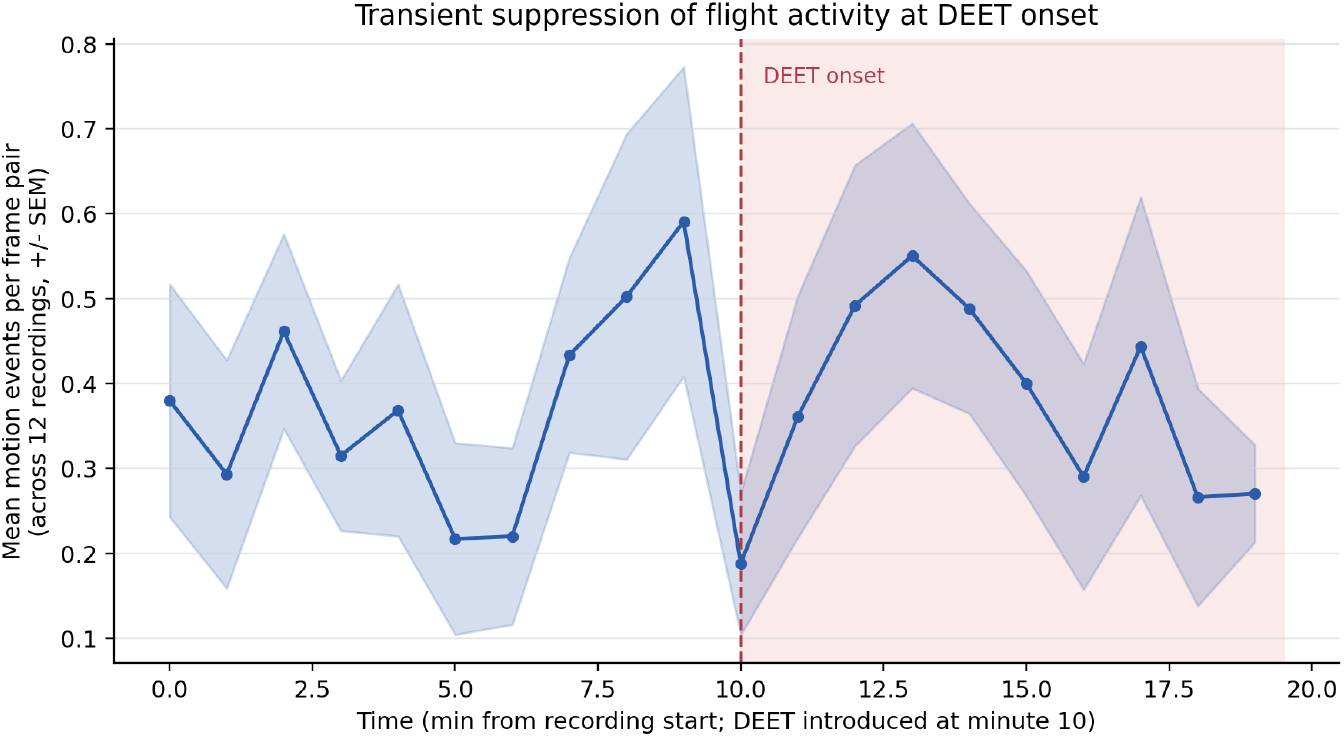
Mean flight motion per frame pair across all 12 recordings, by one-minute bin (mean *±* SEM). DEET is introduced at minute 10 (dashed line; shaded region is the bait-plus-DEET interval). Motion falls sharply at onset and returns to baseline within about three minutes.

### 3.3 Flight drops but the animals stay detectable

The chamber is closed, so the mosquitoes could not leave it and their continued presence is not in itself a finding. What the occupancy readout shows is narrower: the animals stayed detectable in the imaged region while flight collapsed (Figure 2), so the motion drop is not the pipeline losing the animals. Occupancy showed no drop at onset. It declined slowly across the whole recording as the mosquitoes settled, a trend that began well before DEET, and stayed at about 5.5 detections per frame through the DEET half. In the time-matched window of two minutes before and after onset, flight fell from 0.51 to 0.35 events per frame pair (0.19 in the onset minute), while presence changed only from 5.6 to 5.1 detections per frame. Both readouts come from the same frames and region, so the difference lies in behavior, not in the sensor.

**Figure 2:**
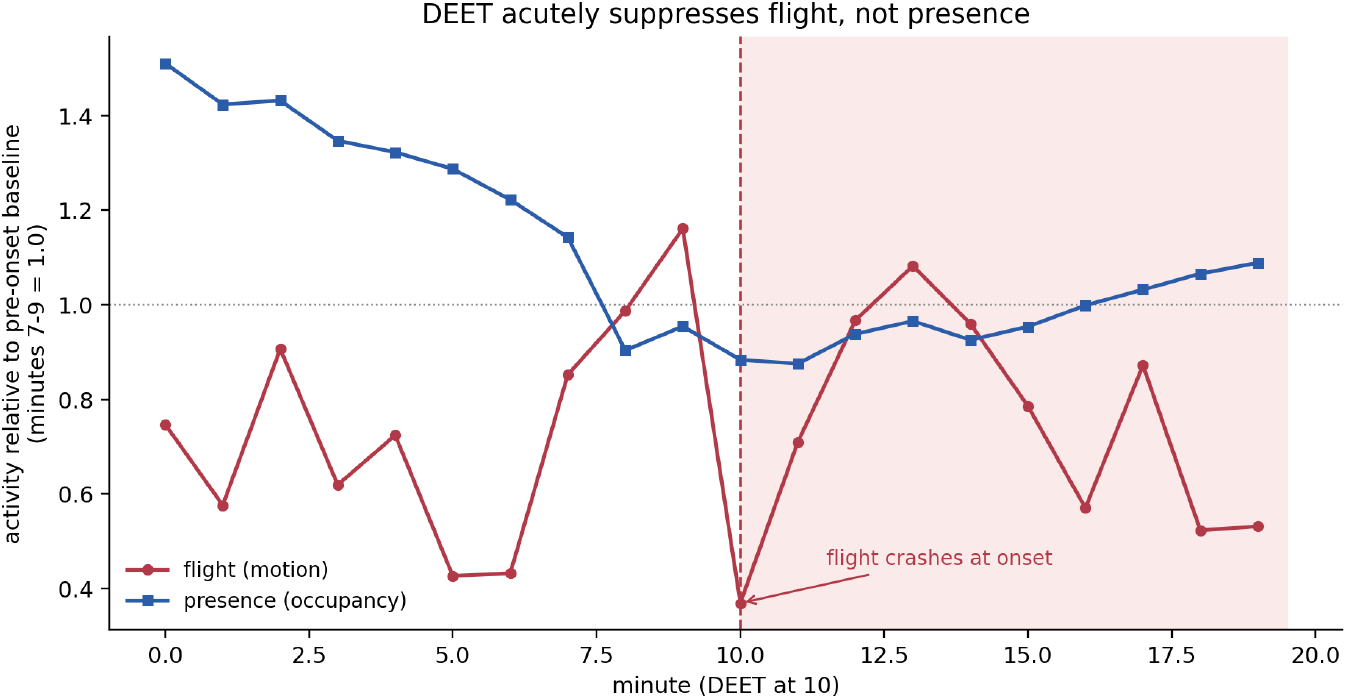
Flight versus presence around DEET onset, pooled across all 12 recordings and normalized to the pre-onset baseline (minutes 7–9 = 1.0). Flight (motion) collapses at onset (dashed line) and recovers; presence (occupancy, counting resting and flying mosquitoes) shows no onset drop. Because the chamber is closed, presence is informative only as a control on the readout: it shows the motion collapse is not the pipeline losing the animals, not that the animals stayed. The slow decline in presence before onset is settling after introduction.

Occupancy alone cannot tell a resting mosquito from a knocked-down one. The time course does: flight returns to near its pre-onset level within about three minutes (Section 3.2), which mortality or lasting knockdown could not produce. We therefore describe the acute response as a transient, reversible suppression of flight. We do not claim it shows non-avoidance; in a closed chamber, movement within the volume is a separate question, addressed only weakly by the zone analysis below. We do not use occupancy to compare colors: because of the settling trend, the full-half occupancy ratio exceeds one for five of six colors whatever their motion direction. Those values are in the released outputs for transparency.

### 3.4 Colors look different at first

The per-color trajectories (Figure S2) and effect sizes (Figure S3; Table 2) suggest that the response differs by color. White had the largest sustained suppression: activity fell by about two thirds at onset (acute drop 0.64), stayed low (late plateau 30% of baseline), and its full-half ratio of means was 2.37 with both sessions above one (5.86 and 2.05). Red had the largest acute drop (0.84), but one session carried its sustained effect (per-session ratios 8.90 versus 1.01), so by magnitude red is suppressive at onset only. By the stated criterion (ratio above one in both sessions) red also passes, on a knife edge: its second-session ratio is 1.0075. Table 2 labels red “acute only” on grounds of magnitude; we flag this because the count of concordant colors matters in Section 3.5. Green, orange, and blue were flat or increased under DEET (full-half ratios *≤* 0.85), and blue’s two sessions disagreed. Yellow was the least active color (pre-onset baseline 0.06 events per frame pair). Its large acute drop and large “recovery” are ratios on near-zero counts and are not interpretable, so we read yellow from its trajectory only. A between-color heterogeneity test was not significant and sat near its null floor (permutation ANOVA *p* = 0.10–0.18; *η*^2^ null 0.45).

**Table 2:** Per-color behavioral fingerprint. All values pool both same-color recordings (ratio of means). Acute drop is the fractional fall in motion in minutes 10–11 relative to minutes 7–9; recovered is the late plateau (minutes 13–19) as a fraction of that baseline; sustained is full bait-only half / full bait-plus-DEET half, with the two per-session ratios for concordance. A value above 1 means less motion under DEET. Yellow^*\**^ has a near-zero baseline, so its acute and recovered ratios are unreliable.

| Color | Baseline | Acute drop | Recovered | Sustained (pooled) | Per session | Concordant |
| --- | --- | --- | --- | --- | --- | --- |
| White | 1.02 | +0.64 | 0.30 | 2.37 | 5.86 / 2.05 | yes (suppressed) |
| Red | 1.05 | +0.84 | 0.36 | 1.64 | 8.90 / 1.01 | acute only |
| Blue | 0.28 | −0.07 | 1.20 | 0.69 | 1.28 / 0.48 | discordant |
| Green | 0.22 | −0.15 | 1.44 | 0.71 | 0.92 / 0.61 | yes (increased) |
| Orange | 0.41 | −0.30 | 0.78 | 0.85 | 0.90 / 0.42 | yes (increased) |
| Yellow* | 0.06 | +0.64 | 10.7 | 0.24 | 0.17 / 0.80 | yes (increased) |

Activity also shifted between zones under DEET, in the direction of avoidance of the source (Table S2). Across colors, the share of activity in the DEET zones (2–3) generally fell. White showed the largest shift toward the far end (activity center of mass +0.64 zones toward the refuge), and its refuge zones (6–7) rose from 10% to 16% of activity. The counts are low, the zone estimates come from the same motion events as the temporal measures, and the order of the shifts broadly follows the baseline activity order. We therefore read the spatial results as consistent with avoidance in aggregate, not as a per-color discriminator.

Taken at face value, this pattern would name white as a candidate color-specific modulator. Section 3.5 shows why it should not be taken at face value.

### 3.5 The color ranking follows baseline activity

The order of the colors by suppression is close to their order by baseline flight (Figure 3A). Across the six colors, baseline flight rate and sustained suppression correlate at Spearman +0.83 (exact permutation *p* = 0.058, *n* = 6); at the recording level the correlation is +0.66 (*n* = 12). Part of this is mechanical, because the baseline rate is the numerator of the suppression ratio, but under the null with no color effect that coupling alone gives a median correlation of only +0.14 (Figure 3B). Most of the association is therefore in the data. That is the problem: color and activity are collinear across six points and cannot be separated.

**Figure 3:**
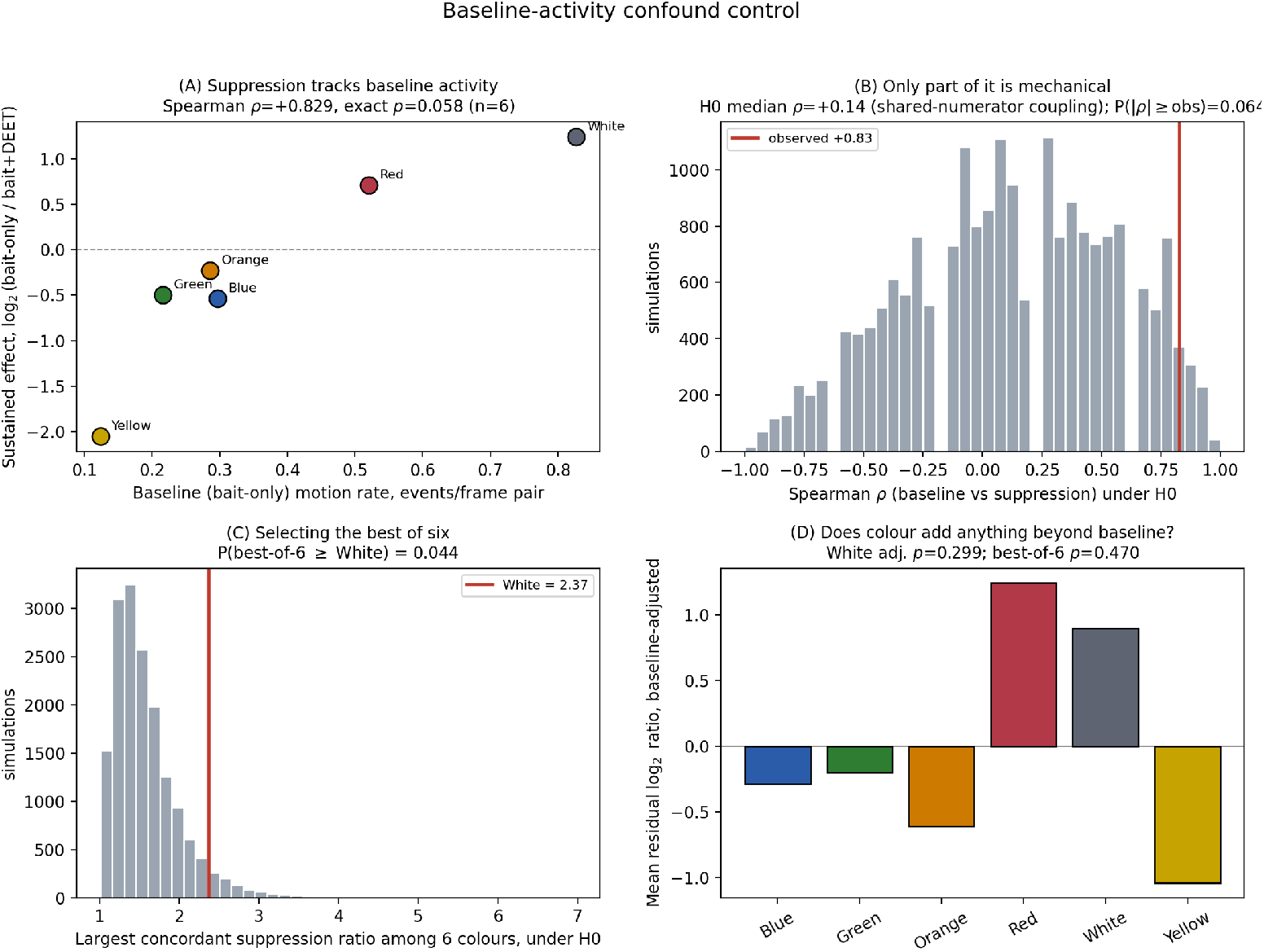
Baseline-activity confound control. **(A)** Sustained suppression against baseline flight rate for each color (Spearman +0.83, exact *p* = 0.058, *n* = 6). **(B)** Null distribution of that correlation when DEET acts identically on every color; the median of +0.14 is the mechanical part from the shared numerator, and the observed value (red line) exceeds it. **(C)** Largest concordant suppression ratio among six colors under the same null, with white’s 2.37 marked; the best of six reaches it in 4.4% of simulations. **(D)** Mean baseline-adjusted residual per color with labels permuted within session; no color is distinguishable (white *p* = 0.30; *p* = 0.47 corrected for best-of-six selection).

Three results follow. First, the concordance criterion does not discriminate. Under the null, a mean of 1.53 of the six colors pass the cross-session concordance bar by chance; we observed two (white and red). With one recording per cell and six colors examined, agreement between two sessions is an ordinary outcome of noise. Second, the magnitude is only mildly unusual. The best of six colors reaches white’s ratio of 2.37 together with concordance in 4.4% of 20,000 null simulations (Figure 3C). Something beyond pure noise may be present, but this test does not attribute it to color. Third, and decisively, color explains nothing once baseline activity is accounted for. Regressing recording-level log_2_ suppression on baseline rate and permuting color labels within session gives *p* = 0.30 for white on its own, and *p* = 0.47 after correcting for having selected the best of six colors (Figure 3D). White’s adjusted residual ranks second of six, behind red. The result is stable across simulation seeds.

The simplest account is that DEET-linked flight suppression scales with how much the animals were flying to begin with, and that the six lighting conditions differed in baseline flight. We cannot exclude a real color effect; absence of evidence at *n* = 2 per color is not evidence of absence. But these data do not support naming any color, and we withdraw that claim. The acute measure does not share this defect: it contrasts adjacent minutes within a recording, so it neither inherits the shared numerator nor rewards high-baseline recordings, and its correlation with baseline activity is *−*0.35. The transient onset result in Section 3.2 is unaffected.

## 4 Discussion

Two findings are robust. In this assay, DEET causes an acute, transient drop in flight: motion roughly halves in the first minute and recovers within about three minutes. It is not a sustained suppression, which fits rapid habituation and the habituation noted in the original thesis [Carver, 2018]. The result is pooled across all 12 recordings, uses a time-matched contrast of adjacent minutes, and does not depend on color, so the confound discussed below does not apply to it. The occupancy readout adds that the animals stayed detectable while flight collapsed. The chamber is closed, so this is not a statement about dispersal; the mosquitoes had nowhere to go, and a claim that DEET failed to drive them away would be unfalsifiable here. Its value is as a control on the readout: a motion-only assay cannot tell animals that stopped flying from animals it stopped seeing, and the occupancy channel separates the two. With the recovery of flight, which rules out mortality, DEET’s acute action here was to ground the animals, not to remove them from view.

The color question we answer in the negative, and we state why. The detectability control shows that the activity differences across colors are behavioral, not a matter of what the camera can see. That control is necessary but not sufficient. A ratio of half-means inherits the baseline it is divided by, and baseline activity was where these lighting conditions differed most. Once baseline is adjusted for and selection among six colors is accounted for, no color stands out. We had first read white’s cross-session agreement as strong evidence; the null simulation shows that about 1.5 colors clear that bar by chance in this design. An analysis that stopped after the first control, as ours initially did, would have reported white as a candidate with apparent caution (effect sizes rather than *p*-values, concordance rather than an omnibus test). None of that restraint would have helped, because the problem was not the strength of the inference but whether the quantity could be identified at all. We think this failure mode is common in small archival reanalyses, where the effect measure is inherited from the original protocol. The visual-integration literature [van Breugel et al., 2015, Vinauger et al., 2019, Alonso San Alberto et al., 2022] makes a color effect plausible but does not predict one, so we leave the question open.

Three limitations bound the interpretation. First, the onset response cannot be fully separated from the physical disturbance of introducing the stimulus. A sudden, recoverable drop is what either a repellent onset or a mechanical disturbance could produce; telling them apart needs a sham (solvent) introduction control, which this archival dataset lacks. We therefore call the response DEET-*linked*, not DEET-*specific*. Second, the design is retrospective and minimally replicated. One recording per color per session means color is crossed with session but not replicated within it, so a formal color-by-DEET interaction cannot be tested, and a real color effect could be present and undetected. Third, the motion readout is an activity proxy, not a count of individuals. The occupancy readout partly addresses this, but it has its own limits. The closed chamber guarantees presence, so presence alone is uninformative. Occupancy cannot separate a resting animal from a knocked-down one; only the recovery of flight does. Persistent resters are removed by the fixed-structure filter. Its full-half comparison is confounded by settling. Neither readout replaces identity-preserving tracking.

The strengths are methodological. All 12 recordings were analyzed with a transparent extension of the thesis pipeline. The ROI was calibrated per session, and incomplete recordings were rejected rather than truncated. Detection was controlled across colors, and the effect measure was audited for confounding. Every statistic reproduces from the deposited tables without the raw images. A prospective study should: (i) include a sham-introduction (solvent) control; (ii) match baseline flight across light conditions or use a time-matched within-recording effect measure, and report per-condition baseline rates; (iii) replicate within condition rather than rely on cross-session concordance, which chance passes in roughly a quarter of colors at this design; (iv) sample the time course densely enough to characterize recovery and dose dependence; (v) record calibrated irradiance for each condition; and (vi) add an occupancy- or trajectory-based readout alongside motion counting.

## 5 Conclusion

We reanalyzed an archived six-color chamber dataset to ask whether the *Aedes aegypti* response to DEET depends on ambient light color. Two findings hold. The DEET response is a short, reversible drop in flight that recovers within about three minutes. The animals stay detectable while flight falls, so the drop is a change in behavior, not a failure of the readout; because the chamber is closed, presence says nothing about dispersal. The color question we answer in the negative: detection is uniform across colors, but the suppression ranking cannot be separated from baseline flight, and no color remains distinct after adjustment and selection correction. We report no candidate color. The lesson carries beyond this dataset: ratio-of-halves suppression measures are confounded with baseline activity, and cross-session agreement is weaker evidence than it looks when several conditions are compared. A prospective study with a disturbance control, within-condition replication, and a time-matched effect measure could still answer the question. These recordings cannot.

## Data and code availability

The analysis pipeline (the `mosquito_movement` Python package) and all machine-readable summaries and figures are released with this manuscript. The principal results derive from `analysis/detectability_control.py`, `analysis/baseline_confound.py`, `analysis/color_fingerprints.py`, and `analysis/occupancy.py` (outputs under `artifacts/`), the full-recording motion pass (`artifacts/baseline_full*`), and the peri-onset pass (`artifacts/baseline_dense_fixed*`). Reproduction steps are in `REPRODUCE.md`. The raw images (about 270,000 frames) are held by the authors and are available on request. The derived per-frame and per-bin tables that reproduce every reported statistic and figure, together with the analysis code, are deposited on Zenodo under DOI 10.5281/zenodo.22239658 (about 9 MB; inventory in `DEPOSIT_MANIFEST.md`; data CC-BY-4.0, code MIT). Every analysis stage after the image reduction runs from the deposited tables alone; with the image drive unmounted, all reanalysis outputs regenerate byte-identically, because the permutation, bootstrap, and simulation procedures are internally seeded.

## Acknowledgments

We thank Brandon J. Carver, whose Master’s thesis built the chamber system, the behavioral paradigm, and the original recordings that this reanalysis rests on.

## Funding

This work received no specific funding.

## Competing interests

The authors declare no competing interests.

## Supplementary material

**Figure S1:**
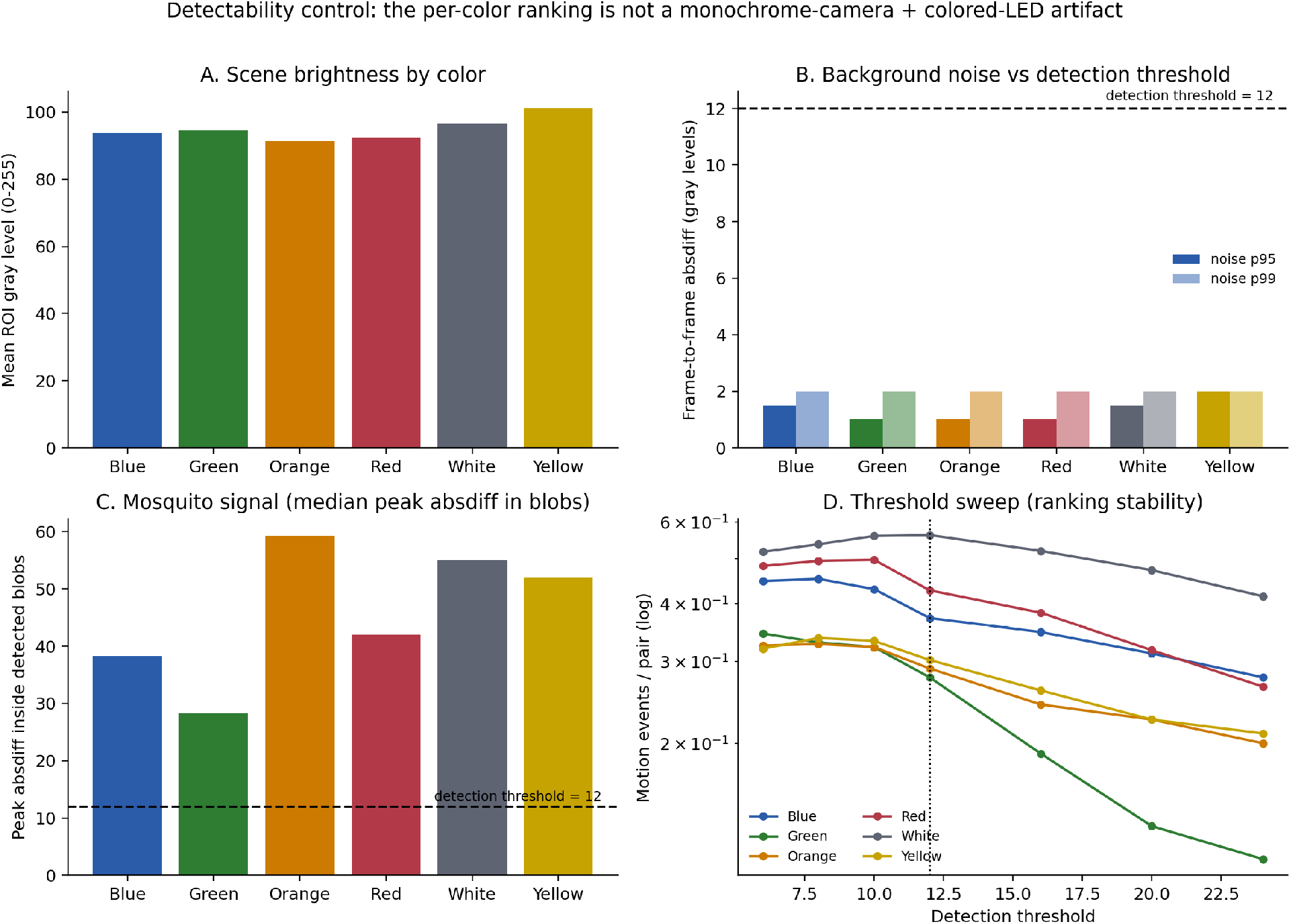
Detectability control. **(A)** Scene brightness by color. **(B)** Background noise floor (95th/99th percentile of mosquito-free frame-to-frame difference) against the detection threshold of 12 (dashed line). **(C)** Median mosquito motion signal, which clears the threshold in every color. **(D)** Threshold sweep: the per-color activity ranking is preserved as the threshold changes.

**Figure S2:**
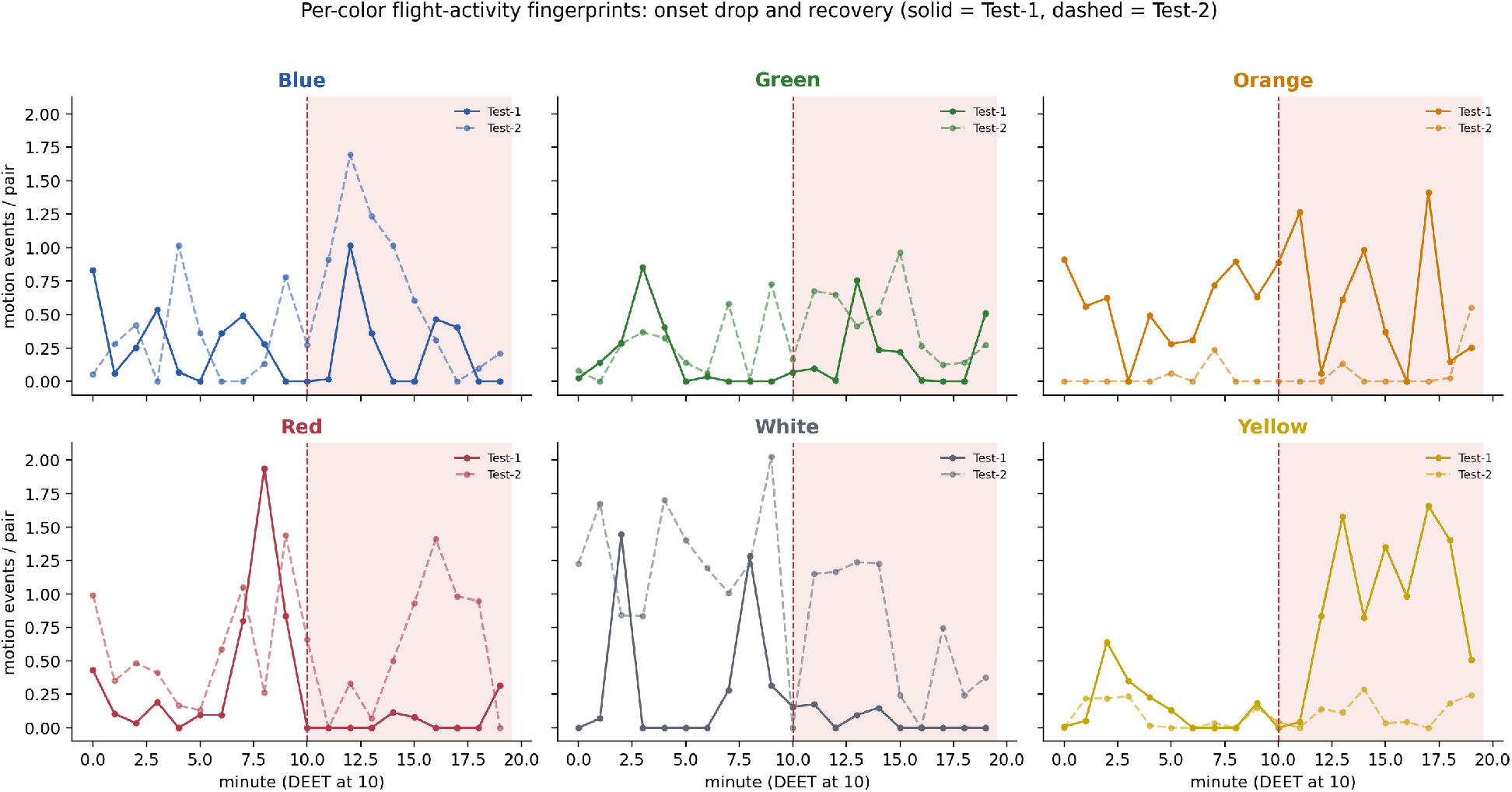
Per-color minute-resolved motion, both sessions overlaid (solid = session 1, dashed = session 2). DEET is introduced at minute 10 (dashed line; shaded = bait-plus-DEET). The onset drop and recovery are clearest for red and white; yellow is sparse throughout the bait-only interval.

**Figure S3:**
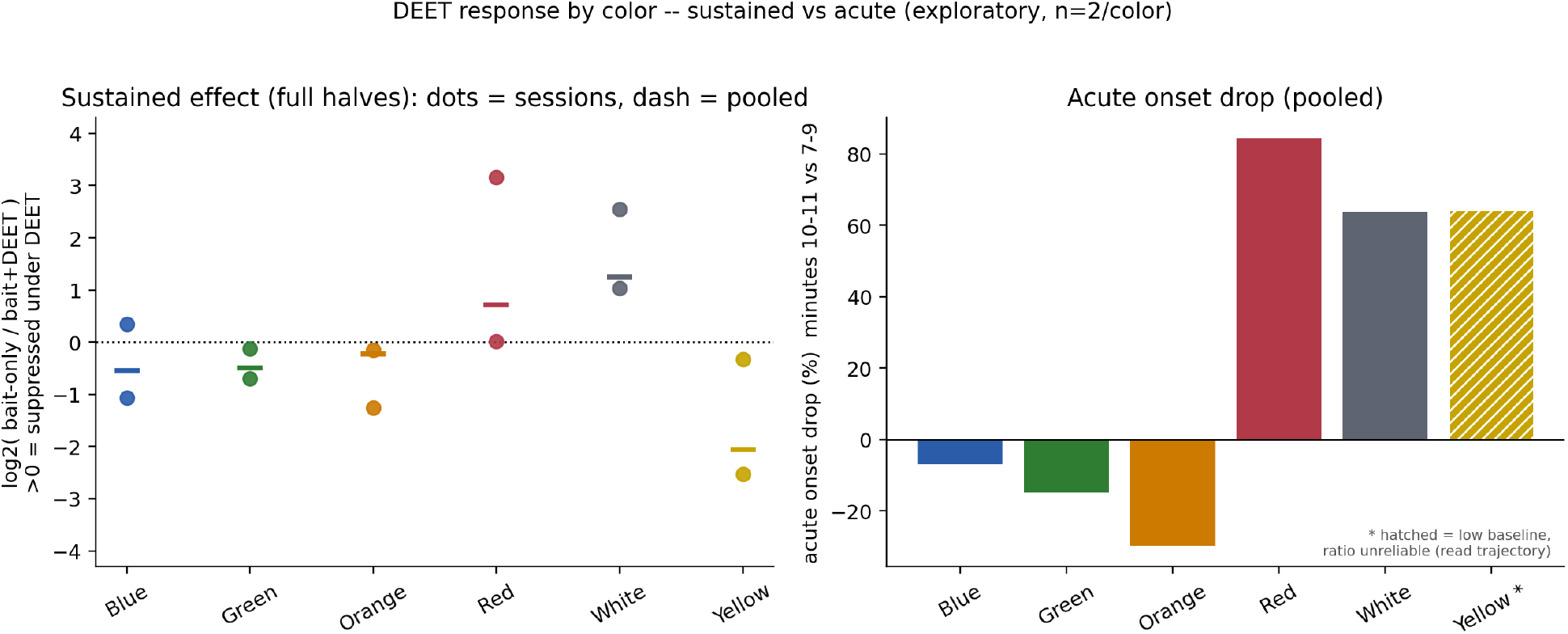
DEET response by color. **Left:** sustained effect over the full halves as log_2_(bait-only / bait-plus-DEET); values above zero mean suppression. Dots are the two sessions; the dash is the pooled estimate. **Right:** pooled acute onset drop (minutes 10–11 versus 7–9); the hatched bar (yellow) has a near-zero baseline and an unreliable ratio. Exploratory, *n* = 2 recordings per color.

**Table S1:** Detectability control, per color (mean of the two sessions). Brightness is the mean ROI gray level (0–255); noise p95 is the 95th percentile of the mosquito-free frame-to-frame absolute difference (gray levels); signal margin is the median peak absolute difference inside detected blobs minus the threshold of 12; SNR is the per-recording signal/noise ratio averaged over the two sessions (a mean of ratios, so it does not reproduce from the printed columns); baseline rate is bait-only motion events per frame pair.

| Color | Brightness | Noise p95 | Signal margin | SNR | Baseline rate |
| --- | --- | --- | --- | --- | --- |
| Blue | 93.7 | 1.5 | +26.2 | 29.2 | 0.30 |
| Green | 94.6 | 1.0 | +16.2 | 28.2 | 0.22 |
| Orange | 91.4 | 1.0 | +47.2 | 59.2 | 0.29 |
| Red | 92.5 | 1.0 | +30.0 | 42.0 | 0.52 |
| White | 96.5 | 1.5 | +43.0 | 42.2 | 0.83 |
| Yellow | 101.1 | 2.0 | +40.0 | 26.0 | 0.12 |

**Table S2:**
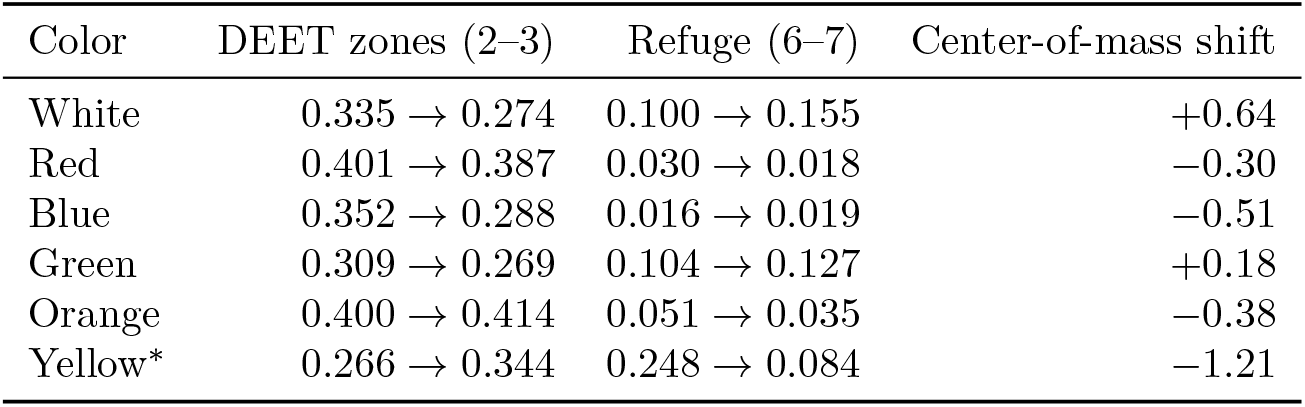
Spatial composition of activity by color (fraction of motion events, bait-only *→* bait-plus-DEET; pooled over sessions). Zones 2–3 hold the DEET source and peak vapor; zones 6–7 are the far, low-DEET refuge. Center-of-mass shift is along the zone 1 (bait) to zone 7 (refuge) axis; positive = toward the refuge. Counts are low; values are suggestive. Yellow^*\**^ is low-baseline.

**Table S3:** Recording-level summary for the sustained (full-halves) contrast. Values are mean motion events per analyzed frame pair; a positive raw difference means less motion after DEET onset.

| ID | Color | Bait only | Bait+DEET | Raw diff. |
| --- | --- | --- | --- | --- |
| Blue-1 | Blue | 0.289 | 0.226 | +0.062 |
| Blue-2 | Blue | 0.304 | 0.635 | −0.331 |
| Green-1 | Green | 0.175 | 0.190 | −0.016 |
| Green-2 | Green | 0.257 | 0.418 | −0.161 |
| Orange-1 | Orange | 0.542 | 0.599 | −0.057 |
| Orange-2 | Orange | 0.030 | 0.071 | −0.041 |
| Red-1 | Red | 0.453 | 0.051 | +0.402 |
| Red-2 | Red | 0.588 | 0.583 | +0.004 |
| White-1 | White | 0.339 | 0.058 | +0.282 |
| White-2 | White | 1.314 | 0.639 | +0.675 |
| Yellow-1 | Yellow | 0.160 | 0.918 | −0.759 |
| Yellow-2 | Yellow | 0.088 | 0.110 | −0.022 |

## Notes

https://doi.org/10.5281/zenodo.22239658

### Competing Interest Statement

The authors have declared no competing interest.

